# Complete loss of mitochondrial complex I genes in mistletoes (Viscaceae) and evidence of polyadenylated mitochondrial transcripts shown by whole transcriptome sequencing

**DOI:** 10.1101/2021.06.11.448138

**Authors:** Athanasios Zervas, Mizuki Takenaka, James Leebens-Mack, Ole Seberg, Gitte Petersen

**Author notes:** **Corresponding authors:** Athanasios Zervas.

## Abstract

The profound absence of mitochondrial complex I (NADH-ubiquinone oxidoreductase) genes from the mitogenome of *Viscum* spp. and the rapid rates of molecular evolution characterizing most of their remaining mitochondrial genes raise questions regarding the possible transfer of the entire *nad* gene set to the nucleus, as well as for the functionality of the remaining highly divergent genes. Using whole transcriptome sequencing in three species of Viscaceae: *V. album*, *V. crassulae*, and *Phoradendron leucarpum* we were able to confirm transcription of all previously identified genes. However, we did not detect any *nad* gene transcripts, thus, providing further evidence of the complete loss of complex I in Viscaceae. The results from transcriptome sequencing also revealed that levels and patterns of RNA editing were not different from those found in autotrophic plant species. Hence, RNA editing is not a means of restoring conserved domains or folding sites of the proteins coded for by the divergent mitochondrial genes. Since we were able to recover mitochondrial genes transcripts following a sequencing protocol targeted towards polyadenylated mRNA molecules, it is suggested that mitochondrial genes undergo post-transcriptional polyadenylation in Viscaceae.

## Introduction

Since the endosymbiosis event of the alpha-proteobacterium 1.5 billion years ago, which gave rise to the mitochondrion that allowed cells to produce the energy they need through the oxidative phosphorylation pathway, the mitochondrial genome has experienced gradual reduction in size and extensive transfer of genes to the nucleus (Lang et al. 1999). The mitochondrial genome includes 5 complexes of genes involved in respiration: complex I (NADH-ubiquinone oxydoreductase), complex II (Succinate dehydrogenase), complex III (Ubiquinol-cytochrome c reducatese), complex IV (Cytochrome c oxidase) and complex V (ATP synthase) (Mower et al. 2012). In animals, the mitogenome has been so reduced that it only harbors essential genes of complexes I-IV, has an average size of 16 kb, very few intergenic sequences, and usually maps to a circular molecule (Boore 1999). In contrast, plant mitochondrial genomes show structural variability and fluidity (Mower et al. 2012), varying from one to many circular molecules (e.g. *Silene conica* L. (Sloan et al. 2012)), to a combination of circular and linear molecules (e.g. *Viscum scurruloideum* Barlow (Skippington et al. 2015)). At the same time, they vary enormously in size, ranging from some 60 kb to over 11 mb, while still only harboring a modest number of relatively conserved genes, but having extended areas of intergenic space including numerous repeats and sequences transferred from the nuclear and plastid genome (Mower et al. 2012). Evolutionary rates of mitochondrial genes depend on the complex or class to which they belong, with genes in complex IV (ATP-synthase genes) evolving faster, and ribosomal protein coding genes evolving slower than the others (Mower et al. 2007). For each complex, certain taxa exhibit increased evolutionary rates, but on average, there is no statistically significant difference between higher plants, with the exception of the Viscaceae (Zervas et al., 2019). Recently, Petersen et al. (2015a) and Skippington et al. (2015) sequenced and assembled the mitochondrial genomes of *Viscum album* L. and *V. scurruloideum*, respectively. The mitogenomes of the two species were very different in structure and size, with *V. album* having a circular mitogenome of 565,5 kb, while *V. scurruloideum* having a much smaller circular mitogenome of 42,2 kb as well as a linear molecule of 23,6 kb. Both groups concluded that *Viscum* mitochondrial genes are very deviating from genes of even closely related taxa, and that they lack all genes of complex I (NAD-ubiquinone oxidoreductase complex). The NADH-ubiquinone oxidoreductase complex (mitochondrial complex I) is the largest in the plant mitogenome (in *A. thaliana* it is comprised of 48 subunits, 9 of which are encoded in its mitogenome (Giegé 2007)). It is there, where the initial electron transfer occurs that starts the oxidative phosphorylation pathway (Remacle et al. 2008). Originally, Petersen et al. (2015) considered additional genes especially in complex IV as lost, but using relaxed annotation criteria and low similarity scores they were recovered by Skippington et al. (2017). Subsequent studies have shown that the heavily modified mitochondrial gene complement may apply to the entire Viscaceae: absence of *nad* genes and increased substitution rate of other mitochondrial genes were found in *Phoradendron liga* (Viscaceae), but other representatives of the Santalales (e.g. *Loranthus* L., Loranthaceae) share the gene content and molecular evolution rates of other angiosperms (Zervas et al. 2019).

The question, however, remains, whether the mitochondrial genes previously identified in *V. album* (Petersen et al. 2015a, Skippington et al. 2017) are in fact expressed, as the only evidence of their existence thus far is *in silico*. And if they are expressed, one may logically speculate that they may be under extensive RNA editing, such that the mature transcript resembles more the “canonical” one from other plants. RNA editing (a post-transcriptional change of specific Us to Cs) was first described in trypanosome mitochondria (Benne et al. 1986), but later found in plant mitochondrial (Covello & Gray 1989) and chloroplast (Hoch et al. 1991) genomes, too. Although most mitochondrial genes include a large number of edited sites (e.g. Giegé & Brennicke 1999), the exact mechanism and enzymes involved in the process are still not clear (Takenaka et al. 2008). To precisely determine which sites are edited, comparison of DNA and RNA sequences is required, but computational tools like the Predictive RNA Editor for Plants (PREP) suite (Mower 2005) can be used to estimate RNA editing *in silico*. However, such tools may be inaccurate when used for non-standard sequences, like the fast-evolving mitochondrial gene sequences of *V. album.*

Another, still unresolved question is whether the missing complex I genes in *V. album* and other Viscaceae have all been transferred to the nucleus, something not yet seen in any other lineage, although the *nad7* gene has been transferred to the nucleus in *Marchantia polymorpha* L. (Kobayashi et al. 1997). The missing genes could possibly be transferred to the nuclear genome but considering the large size of *V. album*’s genome (201 gb (Zonneveld 2010)) it is calculated that 4.1*10^12^ bases output are needed, which translates to 34 High Output, 300 cycles, Illumina NextSeq 500 kits (https://emea.support.illumina.com/downloads/sequencing_coverage_calculator.html) in order to achieve at least 20x genome coverage. Such an approach would increase the costs to produce, store, and analyze the data significantly (Sboner et al. 2011; Muir et al. 2016). Therefore, whole genome sequencing is not the ideal approach to answer the question of a possible transfer of the complex I genes in the nucleus.

An alternative is to target the messenger RNA molecules and perform whole transcriptome sequencing instead. Genes involved in the primary metabolism of any organism are expected to be actively transcribed under standard conditions, without the need of inducing a response (Thellin et al. 1999). Eukaryotic messenger RNA undergoes post-transcriptional modifications with the addition of a poly-A tail in its 3’ end. These tails provide stability to the molecule, facilitate its export to the cytosol, and give the signal to initiate translation (Chekanova & Belostotsky 2003). This feature is unique to nuclear encoded mRNAs, thus organellar (plastid and mitochondrial) transcripts – having a prokaryotic origin – are devoid of such tails. Prokaryotic mRNA molecules show a different type of polyadenylation after transcription, which initiates their immediate degradation by the degradosome after their successful translation to proteins (Hajnsdorf et al. 1995). In a typical growing mammalian cell RNA molecules consist of approximately 80% rRNA, 15% tRNA and 5% mRNA (Lodish et al. 2000). Thus, most transcriptome library preparation kits employ a magnetic oligo-dT beads strategy to purify the meaningful mRNA from the total extracted RNA pool.

Recently, the complete absence of complex I from the mitochondria of *V. album* was investigated using electron microscopy, biochemical assays and proteomics (Maclean et al. 2018; Senkler et al. 2018). Neither of these studies could detect complex I activity and a rearrangement of the respiratory chain was proposed, while the larger part of ATP was suggested to be produced via cytosolic reactions (glycolysis). If this scenario is correct, the *nad* genes could be truly superfluous and lost completely. For the remaining mitochondrial complexes, a decreased activity was also measured in *V. album*, compared to *A. thaliana* (Maclean et al. 2018; Senkler et al. 2018).

In the present study we aim (i) to verify the complete loss of *nad* genes in members of Viscaceae, (ii) to investigate whether the identified mitochondrial genes are transcribed or not, (iii) and if they are transcribed to explore the extent of RNA editing. Finally, (iv) we wish to test the efficiency of PREP-Mt to correctly identify edited sites in *V. album*. Thus, we sequenced the transcriptomes of *V. album, V. crassulae* Eckl. & Zeyh. and *Phoradendron leucarpum* Patsch. to possibly retrieve complex I sequences. We further identified transcripts of other mitochondrial genes present in the three species, and investigated the level of RNA editing by comparing transcribed sequences to previously published DNA sequences (Petersen et al. 2015a, Skippington et al. 2015).

## Materials and Methods

### Collection of plant samples and RNA extraction

Leaves and stems of the same individuals of *V. album* and *V. crassulae* as previously used by Petersen et al. (2015) were collected at the Botanical Garden of the Natural History Museum of Denmark (SNM) in November 2016 and were immediately immersed in liquid nitrogen before being stored at −80°C. Following a similar procedure, leaves and stems of *P. leucarpum* were collected on the campus grounds of the University of Georgia (UGA), U.S.A., in February 2018. Total RNA was extracted from approximately 100 mg of leave and stem tissue of *Viscum* spp. using the Spectrum Total Plant RNA isolation kit (Sigma-Aldrich), at SNM. Total DNA and RNA was extracted from approximately 100 mg of leave and stem tissue of *P. leucarpum* using the DNeasy anr RNeasy Plant Mini Kit (QIAGEN) respectively, at UGA.

### Library preparation and Next Generation Sequencing

For *Viscum* spp. we used the SENSE mRNA Library Preparation Kit for Illumina Sequencing (Lexogen, Vienna), following the manufacturer’s instructions. The protocol uses oligo-dT magnetic beads to target the poly-adenylated RNA molecules of the total extracted RNA, thus removing ribosomal and transfer RNA molecules, as well as messenger RNA molecules of prokaryotic origin. The purified mRNA was quantified in a Qubit 2.0 (Life Technologies) using the RNA assay kit. Illumina adapters were attached as the protocol instructed, after running a qPCR to determine the optimal number of amplification cycles. The finalized libraries were quantified and verified in a Bioanalyzer (Agilent Technologies) before being submitted for Illumina sequencing at the National Sequencing Center, University of Copenhagen, Denmark. Deep transcriptome sequencing was later performed on *V. album* following the same protocol and using a whole HiSeq 4000 lane for Illumina sequencing at the National Sequencing Center, University of Copenhagen, Denmark. The main reasons for choosing *V. album* compared to *V. crassulae* were the availability of the plant in the Botanical Garden, the already sequenced and assembled mitogenome, and the tissue type, which resulted in more successful RNA extractions. Similarly, for *P. leucarpum* we used the KAPPA stranded mRNA-Seq kit (Roche), which follows the same principles as above, and custom Illumina adapters. The finalized library was quantified on a NanoDrop (Thermo-Fisher Scientific) and verified for quality running qPCR in serial dilutions with the specific Illumina adapters used during the library preparation step, and known library standards as reference (KAPA catalogue KK4903) on an Eppendorf Mastercycler EP Realpex^2^. DNA libraries were prepared as described elsewhere (Johnson et al. 2019). Both libraries were submitted for Illumina sequencing at the Georgia Genomics and Bioinformatics Core, University of Georgia, U.S.A. The samples were sequenced with the following chemistries: Initial *V. album* and *V. crassulae* RNAseq: pair-end 251 bp, *V. album* deep RNAseq: single-end 75bp, *P. leucarpum* DNAseq: pair-end 151 bp. *P. leucarpum* RNAseq: 2×81 bp. The difference in chemistry and length was subject to the capacity and queue of the two Sequencing Centers (SNM and UGA) at the different times the samples were submitted for sequencing.

### Data treatment and analysis

The obtained Illumina reads were trimmed for quality and adapters using fastp, under default settings (Chen et al. 2018). The quality controlled sequencing reads were imported in Geneious Prime 2020.2 (https://www.geneious.com). For *P. leucarpum* DNA derived reads, the mitochondrial genes were retrieved following the same approach as described earlier (Zervas et al. 2019). The RNA derived reads were used in map-to-reference runs, using the Geneious mapper and default settings, against a local database comprising mitochondrial complex I gene sequences from *Vitis vinifera* L. (NC_012119). Subsequently, they were mapped against their respective mitochondrial genes (coding sequences – no introns) of *V. album, V. crassulae* (Petersen et al. 2015) and *P. leucarpum* (this study). Consensus sequences were generated in Geneious Prime 2020 using the “majority threshold” and “trim to reference sequence” options.

The mitochondrial gene DNA sequences were imported in PREP-mt (http://prep.unl.edu/cgi-bin/mt-input.pl) for prediction of edited sites under default settings. Briefly, the user enters a protein-coding DNA sequence and selects the codon position of the first nucleotide, as well as the gene the sequence belongs to. The output is a list of the edited sites, as well as the edited gene sequence. These sequences were also imported in Geneious Prime 2020.2.

The DNA, edited RNA (RNAseq) and preditcted edited (PREP-mt) sequences were aligned using MAFFT v7.388 (Katoh et al. 2002) with the following arguments: algorithm FFT-NS-i x1000; scoring matrix 200PAM/k=2; gap open penalty 1.53; offset value 0.123, as implemented in Geneious Prime 2020.2, to investigate the extent of RNA editing.

## Results

From the initial sequencing runs, after quality control, we recovered 13.347.981 paired reads for *V. album*, 3.380.481 paired reads for *V. crassulae*, and 28.399.998 paired reads for *P. leucarpum*. Our analysis was not able to capture any complex I transcripts. We obtained similar results from the second, deep sequencing run of the *V. album* transcriptome, where we recovered, after quality control, 368.102.614 single reads. Surprisingly, we recovered organellar transcripts from all data sets. This is unexpected since we were targeting polyadenylated mRNA molecules, which would be expected to belong to the nuclear genome and not the organellar genomes.

The cumulative data from our analyses, including analyzed genes, predicted and actual edited sites as well as rates of RNA editing in total and in each position are shown in table For *V. album* we retrieved transcripts from all the mitochondrial genes that have been identified in its mitogenome, thus these sequences could also be used for investigating RNA editing. For *V. crassulae* we identified 8 genes, while for *P. leucarpum* we identified 10 mitochondrial genes.

For all 3 species of the analysis, there were clear differences between the predicted edited sites and the experimentally derived RNA edited sites. Thus, in *V. album* out of a total sequence length of 20.326 bp of all gene transcripts, 287 sites were predicted to undergo RNA editing and 236 were identified by sequencing. Between the two number, only 132 sites (46%) were identical, while the 2^nd^ position was favored with 55% of RNA editing occurring there, followed by the 1^st^ (36%) and 3^rd^ (9%), while just 23 events (10%) resulted in silent changes (no change in the amino acid sequence). In *V. crassulae*, out of a total sequence length of 7.639 bp of all gene transcripts, 132 sites were predicted to undergo RNA editing and 94 were identified by sequencing. Between the two number, only 69 sites (52%) were identical, while the 2^nd^ position was favored with 57% of RNA editing occurring there, followed by the 1^st^ (37%) and 3^rd^ (5%), while just 5 events (5%) resulted in silent changes. In *P. leucarpum*) out of a total sequence length of 8.102 bp of all gene transcripts, 90 sites were predicted to undergo RNA editing and 102 were identified by sequencing. Between the two number, only 45 sites (50%) were identical, while the 2^nd^ position was favored with 53% of RNA editing occurring there, followed by the 1^st^ (38%) and 3^rd^ (9%), while just 9 events (9%) resulted in silent changes.

The rates at which RNA editing occurs varies between genes that belong in the same or different complexes. Thus, *atp1* shows the lowest rate in all 3 species in complex IV, followed by *atp8* (data only from *V. album*). Among the shortest in length, *atp9* shows one of the highest rates of RNA editing across all analyzed sequences. In *Viscum* spp. *ccmC* contains the most edited sites per 100bp, followed in *V. album* by *mttB*. In *P. leucarpum, atp9* shows a 3,15% rate of RNA editing. For most of the remaining transcripts analyzed here, there is approximately 1 edited site per 100bp, which is consistent to the total rate of RNA editing sites in all 3 species, namely 1,16%, 1,23% and 1,26% for *V. album, V. crassulae* and *P. leucarpum* respectively. The predicted rate was higher in *Viscum* spp., 1,41% for *V. album* and 1,73% for *V. crassulae*, but lower for *P. leucarpum* with 1,11%, compared to the experimentally calculated rates. For most transcripts, the silent editing occurred on the 3^rd^ position, hence the similar percentages between them. Only exception is *ccmFc* in *V. album* where a silent edited site occurs in position 2 (out of the 15 identified sites). There was found no clear pattern on the intensity of RNA editing sites across the length of the different transcripts, rather certain genes showed a concentration of such sites at the end of the sequence (e.g. *atp1* in all 3 species), or a random distribution (e.g. *cob*). BLASTX searches against NCBI’s nr/nt database showed (when available) that RNA editing did not occur – or was a silent change – on the verified active sites (e.g. in *atp1* in *V. album* – figure 1).

**Figure 1.**
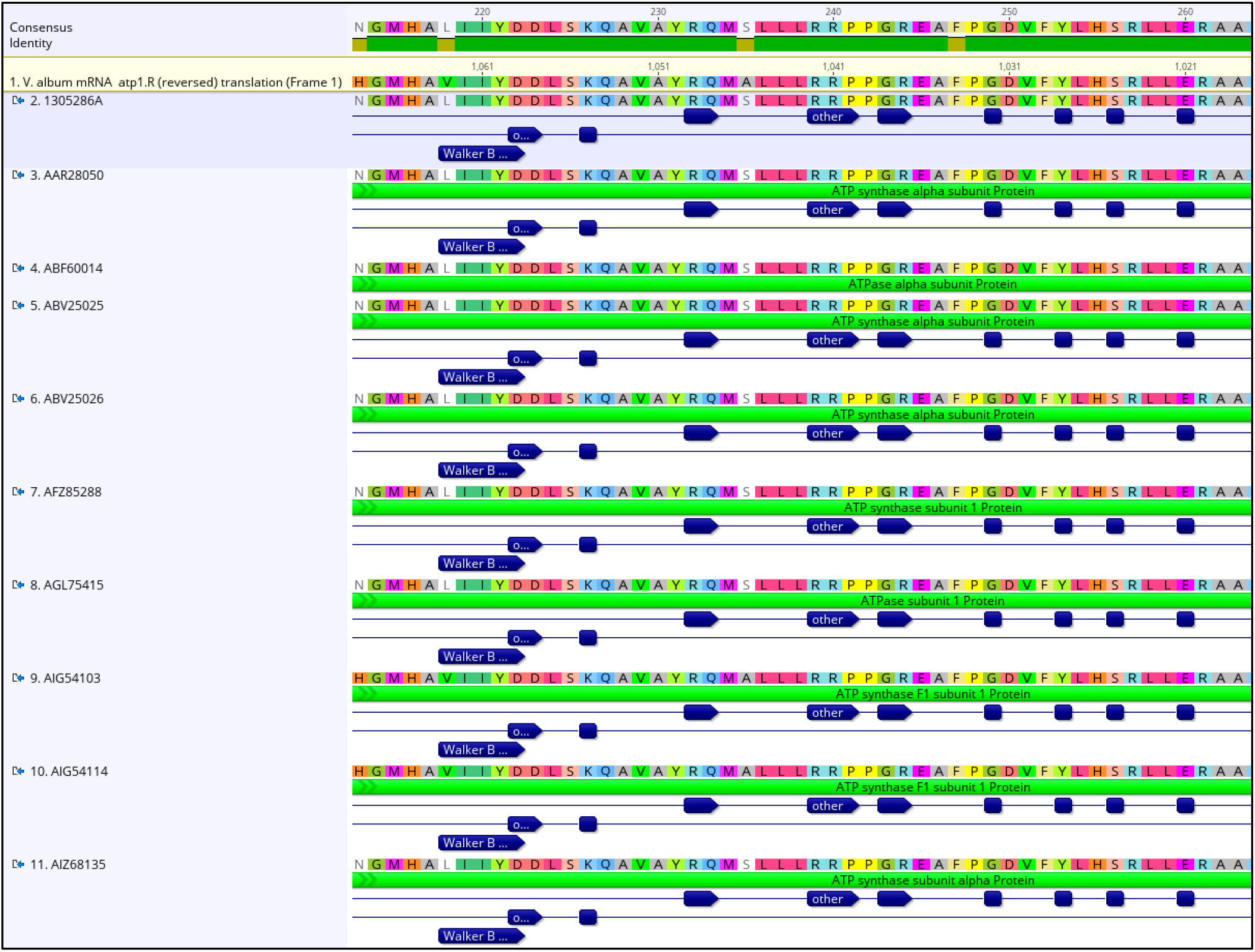
Results from BLASTX analysis of *V. album*’s *atp1* edited transcript. The dark blue boxes and arrows show the verified active site of the gene in the BLAST hits. On the consensus sequence, green denotes no difference between the query and the hits, while yellow boxes indicate a difference in amino acid sequence.

**Table 1:** Targeted genes, Predicted and Actual RNA editing sites and rates of RNA editing in total and in each position for the 3 Viscaceae of the analysis.

| *Viscum album* |  |  |  |  |  |  |  |  |  |
| --- | --- | --- | --- | --- | --- | --- | --- | --- | --- |
| Target gene | Length of sequence (bp) | Predicted Edites sites | RNAseq Edited sites | Identical sites | 1st position | 2nd position | 3rd position | Silent | RNA editing rate |
| atp1 | 1539 | 12 | 7 | 5 | 2 | 5 | 0 | 0 | 0,45% |
| atp4 | 468 | 9 | 6 | 0 | 1 | 4 | 1 | 1 | 1,28% |
| atp6 | 1032 | 19 | 23 | 16 | 9 | 12 | 2 | 2 | 2,23% |
| atp8 | 468 | 2 | 3 | 0 | 2 | 0 | 1 | 1 | 0,64% |
| atp9 | 366 | 7 | 7 | 7 | 1 | 6 | 0 | 0 | 1,91% |
| ccmB | 519 | 32 | 0 | 0 | 0 | 0 | 0 | 0 | 0,00% |
| ccmC | 693 | 24 | 25 | 18 | 9 | 14 | 2 | 2 | 3,61% |
| ccmFc | 2350 | 17 | 30 | 6 | 8 | 15 | 7 | 8 | 1,28% |
| ccmFn | 1782 | 28 | 12 | 7 | 4 | 7 | 1 | 1 | 0,67% |
| cob | 1212 | 11 | 12 | 9 | 9 | 3 | 0 | 0 | 0,99% |
| cox1 | 1833 | 19 | 22 | 18 | 6 | 16 | 0 | 0 | 1,20% |
| cox2 | 753 | 9 | 13 | 9 | 5 | 8 | 0 | 0 | 1,73% |
| cox3 | 741 | 12 | 12 | 10 | 3 | 9 | 0 | 0 | 1,62% |
| matR | 1833 | 12 | 6 | 1 | 1 | 2 | 3 | 3 | 0,33% |
| mttB | 899 | 26 | 29 | 16 | 13 | 14 | 2 | 2 | 3,23% |
| rpl16 | 396 | 1 | 5 | 1 | 0 | 2 | 3 | 3 | 1,26% |
| rps3 | 1549 | 20 | 9 | 1 | 5 | 4 | 0 | 0 | 0,58% |
| rps4 | 972 | 12 | 6 | 3 | 2 | 4 | 0 | 0 | 0,62% |
| rps10 | 276 | 4 | 1 | 0 | 0 | 1 | 0 | 0 | 0,36% |
| rps12 | 420 | 6 | 6 | 4 | 3 | 3 | 0 | 0 | 1,43% |
| sdh4 | 225 | 5 | 2 | 1 | 1 | 1 | 0 | 0 | 0,89% |
| SUM | 20326 | 287 | 236 | 132 | 84 | 130 | 22 | 23 |  |
| PERCENTAGE |  | 1,41% | 1,16% | 46% | 36% | 55% | 9% | 10% |  |

| *Viscum crassulae* |  |  |  |  |  |  |  |  |  |
| --- | --- | --- | --- | --- | --- | --- | --- | --- | --- |
| Target gene | Length of sequence (bp) | Predicted Edites sites | RNAseq Edited sites | Identical sites | 1st position | 2nd position | 3rd position | Silent | RNA editing rate |
| atp1 | 1539 | 12 | 7 | 6 | 2 | 4 | 1 | 1 | 0,45% |
| atp6 | 1245 | 20 | 17 | 12 | 6 | 10 | 1 | 1 | 1,37% |
| atp9 | 234 | 7 | 5 | 5 | 0 | 5 | 0 | 0 | 2,14% |
| ccmC | 693 | 24 | 19 | 9 | 8 | 11 | 0 | 0 | 2,74% |
| ccmFn | 1482 | 37 | 14 | 13 | 2 | 10 | 2 | 2 | 0,94% |
| cob | 1206 | 11 | 12 | 6 | 9 | 3 | 0 | 0 | 1,00% |
| cox2 exon1 | 374 | 7 | 6 | 6 | 3 | 3 | 0 | 0 | 1,60% |
| cox2 exon3 | 68 | 0 | 1 | 0 | 0 | 0 | 1 | 1 | 1,47% |
| cox3 | 798 | 14 | 13 | 12 | 5 | 8 | 0 | 0 | 1,63% |
| SUM | 7639 | 132 | 94 | 69 | 35 | 54 | 5 | 5 |  |
| PERCENTAGE |  | 1,73% | 1,23% | 52% | 37% | 57% | 5% | 5% |  |

| *Phoradendron leucarpum* |  |  |  |  |  |  |  |  |  |
| --- | --- | --- | --- | --- | --- | --- | --- | --- | --- |
| Target gene | Length of sequence (bp) | Predicted Edites sites | RNAseq Edited sites | Identical sites | 1st position | 2nd position | 3rd position | Silent | RNA editing rate |
| atp1 | 1530 | 11 | 9 | 7 | 5 | 4 | 0 | 0 | 0,59% |
| atp6 | 915 | 16 | 22 | 13 | 8 | 13 | 1 | 1 | 2,40% |
| atp9 | 222 | 7 | 7 | 7 | 0 | 7 | 0 | 0 | 3,15% |
| cob | 1224 | 15 | 15 | 9 | 11 | 4 | 0 | 0 | 1,23% |
| cox1 | 1580 | 5 | 18 | 0 | 4 | 6 | 8 | 8 | 1,14% |
| cox3 | 798 | 9 | 8 | 7 | 2 | 6 | 0 | 0 | 1,00% |
| mttB | 782 | 13 | 15 | 1 | 6 | 9 | 0 | 0 | 1,92% |
| rpl16 | 366 | 4 | 4 | 0 | 0 | 4 | 0 | 0 | 1,09% |
| rps10 | 276 | 6 | 2 | 0 | 1 | 1 | 0 | 0 | 0,72% |
| rps12 | 409 | 4 | 2 | 1 | 2 | 0 | 0 | 0 | 0,49% |
| SUM | 8102 | 90 | 102 | 45 | 39 | 54 | 9 | 9 |  |
| PERCENTAGE |  | 1,11% | 1,26% | 50% | 38% | 53% | 9% | 9% |  |

## Discussion

Initially designed to detect transcript of mitochondrial complex I genes potentially transferred to the nuclear genome, we used a standard magnetic oligo-dT bead strategy targeting polyadenylated mRNA transcript. While not recovering any complex I transcript (see below) we most surprisingly discovered transcripts of many other mitochondrial genes. The recovered mitochondrial transcripts, when assembled and aligned to their genomic sequences, showed no signs of degradation. Moreover, the reads map to intron sites as well (i.e. in the entire length of *cox1* – data not shown). Thus, at least in the Viscaceae, plant organellar mRNA molecules either have some type of post-transcriptional modifications similar to nuclear transcripts, or the time-frame between the polyadenylation of the translated transcripts until their degradation by the degradosome (Gagliardi & Leaver 1999; Kudla et al. 1996) is long enough to allow them to be isolated them from total plant RNA pools using the magnetic oligo-dT beads.

The implications of potentially recovering organelle-derived transcripts from a standard magnetic oligo-dT bead approach may be wide. The current approach to studying organellar transcripts (Giegé & Brennicke 1999) including e.g. their RNA editing patterns, is tedious and it is highly dependent on the successful completion of many steps. Firstly, it requires that reference sequences useful for primer design are available. For complete analysis of all the plastid and mitochondrial genes in a given plant, more than 100 primer-pairs would usually need to be designed, and these primer pairs may not be useful for other studies if the target sequences are slightly different. Preferably, the primer pairs also need to have similar properties allowing them to be run in parallel PCRs. Finally, the resulting PCR products need to be verified for length, purified and Sanger sequenced. Thus, the alternative approach is significantly time saving, and whole genome and transcriptome sequencing may even be done in the same run. However, it remains to be determined whether the approach can be applied to other plants, or whether members of the Viscaceae are also unique in terms of their mitochondrial transcription.

The present study demonstrates that the rapidly evolving mitochondrial genes of the three Viscaceae examined here are actively transcribed, despite how divergent they are compared to other angiosperms (Petersen et al. 2015a, Zervas et al., in review). Even the most divergent genes initially considered missing in *V. album* by Petersen et al. (2015a), but subsequently identified by Skippington et al. (2017) are transcribed. Also, two genes, the maturase gene *matR* and the cytochrome C maturation protein subunit B gene *ccmB*, which by Skippington et al. (2017) were proposed to be acquired through horizontal transfer and thus not likely to be functional, were recovered from the transcripts. Comparison between the transcribed sequences and the DNA sequences further reveals that both of the genes experience RNA editing, albeit only one site is edited in *ccmB*. Transcription and editing of the genes suggest that they may both be functional although this needs verification. But whether the genes were actually acquired through horizontal transfer also needs verification possibly by phylogenetic analysis using a deeper taxonomic sampling.

Our analyses of RNA editing show that editing is not more frequent in any of the investigated species compared to normal level in autotrophic plants (e.g. Giegé & Brennicke 1999). Further, the edited sites are not primarily in key positions, where a change in the nucleotide sequence leads to a change in the amino acid that would correspond to the specific triplet and a “proper” residue of an active site. In order for a protein to be functional, both correct folding and the right amino acids in the active or binding sites are a vital requirement. The translation of the edited transcripts still reveals great differences when aligned to canonical protein sequences from other angiosperms. This difference in nucleotide sequence is likely to affect the structure of the resulting product, and possibly also protein activity. Thus, the highly divergent sequences may explain the decreased enzymatic activity observed in the recent biochemical and proteomics assays from *V. album* (Maclean et al. 2018; Senkler et al. 2018).

Testing the efficiency of PREP-Mt (Mower 2005) to predict edited sites revealed a remarkable poor performance. PREP-mt predicted correctly only 58.5% and 67% of the experimentally verified sites in *V. album* and *V. crassulae* respectively, while 41% and 32% of the edited sites were overlooked and 55% and 52% sites erroneously assigned in *V. album* and *V. crassulae* respectively. Previous studies have demonstrated that PREP-Mt is rather accurate for standard mitochondrial genes (e.g., Chaw et al. 2008; Cuenca et al. 2010) although another study found a 10% error rate (Wang et al. 2012). The main cause of its inefficiency in this study is most likely the very divergent sequences of the specific taxa.

In addition to divergent mitochondrial genes, the absence of complex I genes from the mitogenome of the Viscaceae has been demonstrated with the release of two complete *Viscum* mitogenomes (Petersen et al. 2015a; Skippington et al. 2015), and from investigations including 2 other *Viscum* species (Petersen et al. 2015a) and the Argentinian mistletoe *Phoradendron liga* (Zervas et al. 2019). Initiated prior to the recently published additional evidence of complete lack of the complex I genes from the genome of *V. album* (Maclean et al. 2018; Senkler et al. 2018), one of the aims of the current study was to test the potential presence of the genes in the nucleus. Using the approach targeting nuclear polyadenylated mRNA transcripts, we did, however, not detect any complex I transcripts, providing further evidence of their complete absence from the genome. Following the functional transfer to the nucleus of *cox2* in legumes (Adams et al. 1999) and *nad7* in *Marchantia* (Kobayashi et al. 1997) it would not have been entirely surprising for mistletoes to have followed a similar route. However, all current results point towards a complete absence of complex I. Normally, electrons enter the oxidative phosphorylation pathway in mitochondria through the activity of NADH-ubiquinone oxidoreductase, which removes a proton from NADH and transfers the freed electron to ubiquinone (Rich & Maréchal 2010). Petersen et al. (2015a) speculated on other possible pathways *Viscum* spp. might have taken to solve this predicament, one of which was the recruitment of alternative NAD(P)H dehydrogenases. Recently, a biochemical and proteomics study aiming at defining the oxidative phosphorylation system in *V. album* was published (Senkler et al. 2018). A combination of blue native PAGE electrophoresis, coupled to Mass Spectrometry led to the identification of complexes II, III, IV, V and cytochrome c, as well as supercomplexes of complexes III, IV and cytochrome c. Complex I was completely missing and at the same time the relative abundance of complex V (ATP complex) was quite low. However, NAD(P)H and alternative oxidases were recovered, as was hypothesized. The authors recreated an alternative oxidative phosphorylation pathway for *V. album* which is simpler than the canonical one found in other flowering plants (e.g. *Arabidopsis thaliana, Solanum tuberosum*). At the same time, Maclean et al. (2018) performing a similar approach, also showed – biochemically – the complete absence of complex I proteins from mitochondrial extracts. They proceeded with proteome analysis using liquid chromatography-tandem mass spectrometry, from which they isolated 292 proteins that encompass 193 experimentally confirmed in *A. thaliana* mitochondrial proteins. Searching for Gene Ontology terms and comparing them to *A. thaliana*, no significant differences were shown for basic mitochondrial functions, like vitamin biosynthesis, tri-carboxylic acid cycle and mitochondrial organization. However, their relative abundance in *Viscum* relative to *Arabidopsis* was much lower. These results were coupled to ^14^C-glucose isotope flux analysis, which showed a redirection of the carbon metabolic fluxes from the TCA cycle to glycolysis as means of ATP production. This strategy, however, comes at a great metabolic cost. One glucose molecule can produce up to 38 ATP equivalents (usually a net gain of 30 ATP per glucose) through the TCA cycle (Berg et al. 2002), while the net gain from glycolysis is only 2 ATP equivalents. This means that ATP production through glycolysis requires 15 times more glucose compared to aerobic respiration in the mitochondria. It is quite unlikely that the Viscaceae can produce that much more glucose from photosynthesis to cover their energy requirements. The high transpiration rates of the *Viscum* seedling after its establishment on the host (Escher et al. 2008) have suggested a significant carbon flux from the host to the parasite that would sustain the increased metabolism (Těšitel et al. 2010). However, the recent findings may suggest that this transfer of nutrients decreases over time, as the enzymatic activities measured in *Viscum* were lower compared to *Arabidopsis* (Maclean et al. 2018). The low growth rates of the mature Viscaceae could also reflect the presence of polyadenylated transcripts of organellar origin. In plant mitochondria, ribonucleases readily degrade polyadenylated mRNA as means to regulate the levels of the different products (Gagliardi et al.). Thus, low growth rates due to insufficient ATP could result in a reduced enzymatic activity of those ribonucleases, which in turn may not be able to degrade the polyadenylated mRNA fast enough. This could also possibly explain why we were able to retrieve organellar transcripts from the total RNA pool.

In summary, in the present study we set out to investigate the presence of mitochondrial complex I transcripts in *V. album, V. crassulae* and *P. leucarpum*, and to verify active transcription of the remaining highly divergent mitochondrial genes. Following whole transcriptome sequencing we were not able to identify any of the complex I genes, thus providing further molecular evidence of their absence. By designing primers targeting specific mitochondrial gene products in *V. album* we verified their transcription, but also whole transcriptome sequencing recovered mitochondrial gene transcripts suggesting that, at least in the Viscaceae, organellar messenger RNAs undergo a type of polyadenylation. Further analyses of RNA editing in the retrieved transcripts showed that levels of editing are normal and cannot restore amino acid similarity to the active or binding sites that we were able to investigate. Our results complement the recent biochemical and proteomics studies conducted in *V. album* that show loss of the complex I, reduced enzymatic activity of the remaining complexes, and propose a different mechanism of producing ATP involving glycolysis as the main pathway (Maclean et al. 2018; Senkler et al. 2018), as was hypothesized initially (Petersen et al. 2015a).

## Acknowledgements

Special thanks to Charlotte Hansen (SNM) and Amanda Lynn Cummings (UGA) for capable laboratory assistance. The authors thank the staff at the National High Throughput DNA Sequencing Centre of the University of Copenhagen and the staff at the Georgia Genomics and Bioinformatics Core of the University of Georgia for Next Generation Sequencing.

## Data availability

All sequences are available and will be uploaded to GenBank.

## Author’s contribution

AZ, GP, MT and OS conceived and designed the study. AZ carried out the laboratory work, data treatment, data analysis, and drafted the manuscript. GP and OS helped drafting the manuscript. MT, JLM, GP and OS critically revised the manuscript for important intellectual content. All authors read and approved the manuscript.

